# The Expression of Inflammatory Genes in 3T3-L1 Adipocytes Exhibits a Memory to Stimulation by Macrophage Secretions

**DOI:** 10.1101/336610

**Authors:** Dale Hancock, Luxi Meng, Mira Holliday, Nicole Yuwono, Ning Zhang, Gareth Denyer

**Affiliations:** School of Life and Environmental Sciences, University of Sydney, NSW 2006, Australia

**Keywords:** Adipocytes, Inflammation, Macrophages, Transcriptional Memory

## Abstract

Obesity is characterized by increased output of inflammatory compounds from adipose tissue. Whilst the relative contribution of adipocytes and resident macrophages to this phenomenon is debated, there is no doubt that the secretions of each cell type can stimulate the expression of inflammatory genes in the other. We hypothesized that mechanisms must exist to prevent an escalating positive feedback loop between the two cell types, so that after an initial exposure to macrophage secretions, adipocytes would become desensitized to subsequent inflammatory stimulation.

We used microarrays to investigate the response of 3T3-L1 adipocytes to macrophage secretions (macrophage conditioned medium, MCM). MCM caused a rapid (<4 hours) and high amplitude (over 100-fold) rise in the expression of several inflammatory genes. For some genes, generally cytokines, expression returned to basal levels within 24 h following removal of the MCM, but other transcripts, notably those for acute phase proteins and extracellular matrix remodeling proteins, remained highly expressed even during the washout period.

Unexpectedly, some cytokine genes (e.g., iNOS, IL-6) showed an enhanced expression to a second exposure of MCM, illustrating that the transcriptome response of 3T3-L1 adipocytes retains a memory to the first stimulus. We characterized the parameters that give rise to the memory phenomenon, finding that additional stimuli do not augment or abrogate the effect. The memory is preserved for several days after the initial exposure and it is not due to a change in sensitivity to the MCM but, rather, a change in the capacity of the signal-target system. The possible mechanisms of the memory are discussed, along with the physiological ramifications should the phenomenon be replicated *in vivo.*

## INTRODUCTION

Two decades ago, the mechanistic relationship between elevated adipose tissue mass and obesity-related disease was unclear. It is now known that, in the obese state, adipose tissue secretes a wide range of inflammatory mediators, and that elevated systemic levels of these compounds contribute to both Type 2 Diabetes and cardiovascular disease, as recently reviewed by (1). A more complete understanding of how the production of inflammatory agents from adipose tissue is regulated is likely to lead to better therapeutic strategies for the treatment of obesity related disease.

Many different types of inflammatory secretions emanate from obese adipose tissue, and cytokines comprise one of the major groups. Both adipocytes and the macrophages resident in adipose tissue are potential sources of these cytokines and there is some controversy regarding which of these two cell types is mainly responsible for the release of most of the inflammatory hormones. This issue is especially difficult to unravel since, in the obese state, adipose tissue becomes infiltrated with macrophages (2)

The situation is further complicated by the fact that agents released from adipocytes can activate macrophages and *vice versa*. Incubation of adipocytes with the culture medium collected from M1-macrophages (macrophage conditioned medium, MCM) causes a rapid and high-amplitude rise in inflammatory gene expression in both human and mouse adipocyte cell lines, and that this rise is accompanied by increased secretion of cytokines (3) (4) (5) (6) and at least one of these cytokines (TNFα) independently induces similar responses (7) (8) (9). ‘Inflamed’ adipocytes not only secrete agonists that increase the expression and production of cytokines in macrophages, they even express proteins (like MCP-1) which attract even more macrophages (10). Clearly, these interactions have the potential to form a positive feedback loop that could exacerbate systemic inflammation.

Microarray studies have revealed that the response of adipocytes to MCM does not just involve the expression of cytokine mRNAs but is also characterized by rapid and sizeable rises in the levels of extra-cellular matrix remodeling transcripts and repression of adipocyte-specific genes (11) (12). Interestingly, despite the fact that these rises in mRNA levels are often >100-fold over basal, the changes are generally transient, even if the stimulatory milieu is maintained. This waxing and waning of gene expression is a common property in immune cells in response to inflammatory stimuli, and is caused by compensatory and counter-regulatory responses, with prolonged or prior exposure of many types of cells to hormones results in a dampening and loss of sensitivity to subsequent exposures. This is due, in part, to decreased receptor levels (13) (14) (15) and persistence of the post-stimulation dampening responses (16) (17).

Since adipocytes exhibit this kind of response to insulin, showing reduced insulin sensitivity after exposure to high insulin concentrations (13) (14) (18), we reasoned that fat cells might show a similar repression to further inflammatory stimuli after prior exposure to macrophage secretions. It is intuitive that such a mechanism should exist to prevent a rampant positive feedback loop between adipocytes and macrophages. In sharp contrast to these expectations, we describe here experiments that demonstrate quite the reverse: namely, that exposure to inflammatory stimuli leaves an imprint on fat cells that actually renders them more responsive to subsequent stimulation by macrophage secretions.

## METHODS

### 3T3-L1 Cell Culture

Low passage number 3T3-L1 cells were cultured in DMEM (Life Technologies, Australia) which contained 25 mM glucose, 4 mM L-glutamine and 1 mM sodium pyruvate supplemented with 10% (v/v) fetal bovine serum (referred to as DMEM/FBS). Cells were cultured in 6 or 12 well plates in a humidified incubator at 37°C supplemented with 5% (v/v) CO_2_.

Two days post-confluence, differentiation was induced with a cocktail of stimulants; 2 μg mL^−1^ insulin, 0.5 mM isobutylmethylxanthine, 0.25 μM dexamethasone and 2 μM rosiglitazone (all purchased from Sigma) in DMEM/FBS. After three days, the differentiation medium was removed and cells were maintained in post-differentiation medium (DMEM/FBS with 2 μg mL^−1^ insulin) for a further 6–9 days, with media changes every two days or as required. Differentiation was monitored both by microscopy and Oil-Red O staining.

### Production of MCM

The murine monocyte cell line, RAW264.7 was employed to obtain macrophage-conditioned medium (MCM). Cells were cultured until confluence in DMEM/FBS in a CO_2_ supplemented humidified incubator at 37°C. To stimulate an inflammatory response, the cells were treated with lipopolysaccharide (LPS, 50 ng/mL) for 4 h. At the end of this time the incubation medium was collected and filtered through a 0.22 μm filter (Millipore) to remove any debris and cells. The MCM was stored at −80°C until required. Any variations from this procedure are described in the relevant figure or table legend.

### Memory Experiment setup

For the initial stimulation, 9-day differentiated 3T3-L1 cells were treated for 4 h with media containing MCM diluted 1:1 with DMEM/FBS. Following this, the medium was removed, and the cells were washed with PBS and then fresh DMEM/FBS was added to the wells for a further 24 h “washout” period. The medium was then removed and these cells were exposed to a second challenge with MCM diluted 1:10 with DMEM/FBS for 4 h. Total RNA was isolated from cells at 0, 1, 2 and 4 h after the addition of the second MCM challenge (28, 29, 30 and 32 h after the initiation of the first MCM stimulation).

Control cells, which had only been exposed to MCM for one challenge, were also prepared. These were age matched (10-day differentiated) 3T3-L1 cells from the same batch and passage number as above, and there were incubated with MCM diluted 1:10 with DMEM/FBS for 4 h. Total RNA was isolated at 0, 1, 2 and 4 h after the addition of the MCM. Any variations to this scheme are described in the relevant figure or table legend.

### RNA Isolation

Total RNA was isolated by Trizol (Sigma-Aldrich, Australia), using a variation of the method of Chomczynski and Sacchi (19). Essentially the media was removed from each well and the adherent cells were washed once with PBS followed by lysis in Trizol reagent. The lysate was then extracted with chloroform (5: 1 vol:vol) at room temperature. Following centrifugation at 12,000 × *g* for 15 min the upper aqueous phase containing the RNA was removed and the RNA precipitated overnight at 20°C with 1.25 volumes of isopropanol. The pellet, collected after centrifugation at 12,000 x *g* for 30 min, was washed twice with 75% (v/v) ethanol then re-suspended in nuclease-free water. The yield and purity of the RNA was estimated by UV spectrophotometry. The integrity of the RNA was further assessed by denaturing agarose gel electrophoresis (1% (w/v) agarose in 2.2 M formaldehyde MOPS) (20) followed by ethidium bromide staining and visualization using a UV trans-illuminator.

### Microarray analysis

Microarray analysis was performed on RNA samples in order to get a complete perspective of gene expression changes after MCM stimulation. Isolated RNA was treated for application onto Affymetrix GeneChip Gene Mouse 2.0 ST arrays by the Ramacotti Centre at the University of NSW, according to the manufacturers instructions. Microarray data was analyzed using an in-house designed package as has been employed in several other studies (21) (22) (23) (24) (25) (26) (27) (28) (29) (30).

### cDNA synthesis and qPCR

Complementary DNA from each RNA preparation was synthesized with Reverse Transcriptase (Bioline, Australia) according to the manufacturer’s protocol. 500 ng samples of total RNA were primed using random hexamers in 20 μL reaction mixtures containing 500 μM dNTP, 50 mM Tris HCl pH 8.6, 40 mM KCl, 5 mM MgCl_2_, 1 mM MnSO_4_, 1 mM DTT and 0.5 U/μL RNase inhibitor. Primers were annealed at 25°C for 10 min, cDNA synthesis was then carried out at 45°C for 30 min with 10 U/ μL reverse transcriptase followed by enzyme inactivation at 85°C for 5 min.

qPCR was performed in 20 μL reactions using SybrGreen (FAST SybrGreen master mix, Applied Biosystems, Australia) amplifying two different amounts of each cDNA preparation (10 ng/ reaction and 1.25 ng/ reaction) for each primer set. A two-step thermocycler program was employed using an Applied Biosystems 7500 Fast instrument. This was 10 min at 50 °C pre-incubation to remove any contaminating amplified product from previous PCR assays, then 95°C for 20 s followed by 40 cycles of 3 s at 95 °C then 30 s at 60 °C. A melt curve was performed on all reaction products. and each *Ct* normalized to its respective 18S value. As a *Ct* difference between the 10 ng reaction value and its corresponding 1:8 (1.25 ng) value of 3 indicates a quantitative, consistent amplification only samples which showed this *Ct* difference between dilutions were used in subsequent analysis. Primers were designed using a combination of the UCSC Genome browser (https://genome.ucsc.edu/) and Primer3 Plus (www.bioinformatics.nl/primer3plus) with at least one primer in each pair hybridizing across an exon-splice sit, thus restricting amplification products to processed transcripts.

## RESULTS

### A subset of adipocyte-expressed genes display a memory effect after the first of two challenges with MCM

To assess the effect of MCM on adipocytes in culture, we exposed mature 3T3-L1 cells to MCM for up to 24 hours, either with or without a washout of the stimulus at 4 h. Figure 1 shows that the transcriptional output of inflammatory targets in 3T3-L1 adipocytes responds to macrophage secretions in the same way as reported by others (31) (3) (11). The MCM caused a rapid (within 2 h) and high amplitude (>100-fold) rise in the levels of both IL-6 and iNOS transcripts, two key indicators of the adipocyte inflammatory response. Moreover, despite the continued presence of the MCM stimulus, the level of both these mRNAs decreased after 4 h of exposure. In neither case did transcript levels return to basal values, even after a further 24 h incubation. In contrast, when the MCM stimulus was removed and the cells incubated in fresh culture medium, the expression of both genes declined to pre-exposure levels within 8–12 h.

**Figure 1.**
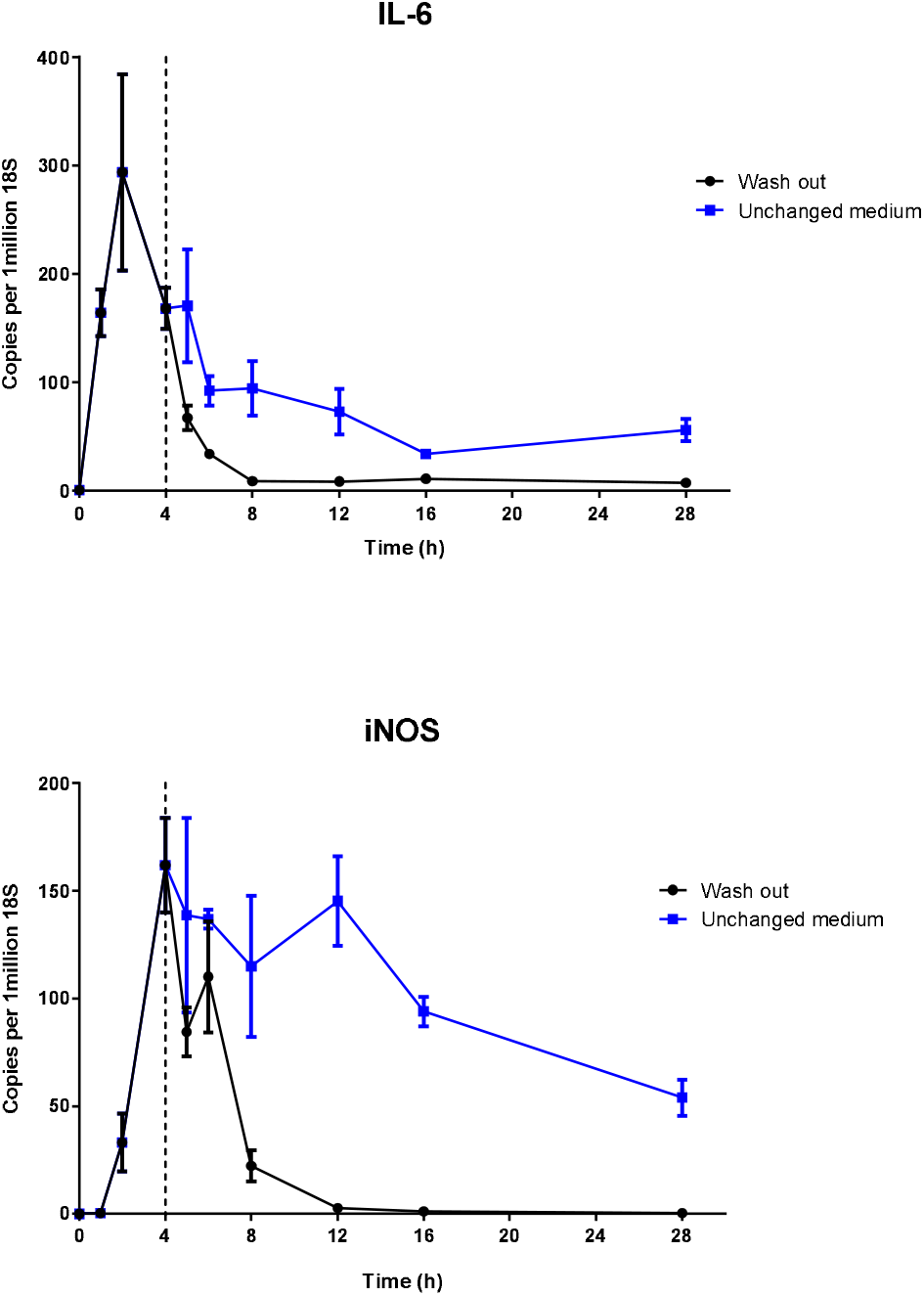
Expression of inflammatory genes in 3T3-L1 adipocytes that received a 4h MCM stimulation followed by a washout or continuous stimulation. Mature 3T3-L1 adipocytes were initially stimulated with MCM for 4h. Medium was either replaced with fresh normal growth medium (washout •) or was unchanged (unchanged medium ■) for 24h. cDNA was synthesised from RNA extracts and gene expression levels were analysed using qPCR. C_T_ results were normalised to 18S rRNA and expressed as transcript copies per 10^6^ 18S copies. Data are shown as mean ±SEM (*n*=3). Statistical analysis was performed using students t-test, p< 0.05*, <0.01**, <0.001***

Although washout of the MCM stimulus appears to reverse the effects of the inflammatory exposure (at least with respect to iNOS and IL-6 expression), we were curious to know if the previously-exposed cells would behave differently to naive cells when exposed to MCM again. Figure 2 shows that cells previously exposed to MCM show an enhanced response to a second challenge. These results indicate that the first exposure to MCM leaves some imprint on the gene expression apparatus of iNOS and IL-6 in adipocytes.

**Figure 2.**
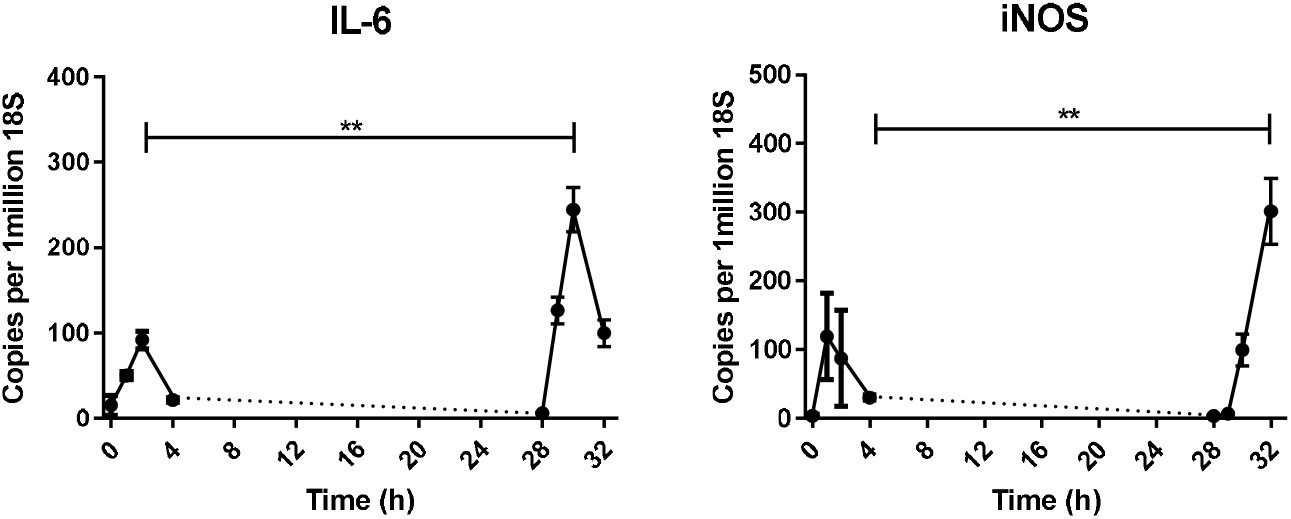
Expression of genes in 3T3-L1 adipocytes during primary and secondary MCM exposures. 3T3-L1 adipocytes were exposed to MCM for 4h (primary stimulation) and then exposed again for another 4h (secondary stimulation) after receiving a 24h washout in normal complete growth medium. cDNA was synthesised from RNA extracts and gene expression levels were analysed using qPCR. C_T_ results were normalised to 18S rRNA and expressed as transcript copies per 10^6^ 18S copies. Data are shown as mean ±SEM (*n*=3). Statistical analysis was performed using students t-test, p< 0.05*, <0.01**, <0.001***, <0.0001****

We next asked whether other transcripts also exhibit this transcriptional memory. We performed microarrays on mRNA isolated from cells exposed to MCM for 0–4 hours, either as a primary stimulation (i.e., cells that had never seen MCM before) or as a secondary stimulation (i.e., after a previous 4-hour exposure followed by a 24-hour washout). As observed by others (11) (31), the expression of several hundred genes was stimulated after MCM exposure (original data available in NCBI Gene Expression Omnibus). Of these, about 20 targets show a greater response to MCM after a previous exposure. Table 1 lists the genes that exhibited: a) a rise in response to the first MCM exposure of >4-fold at least one time point, b) a return to basal (or near-basal) levels of expression after washout, and c) a secondary response to MCM that was at least 2-fold greater than the first exposure at least one time point. In every case, the targets listed in Table 1 have at least one inflammatory ontological classification.

**Table 1.**
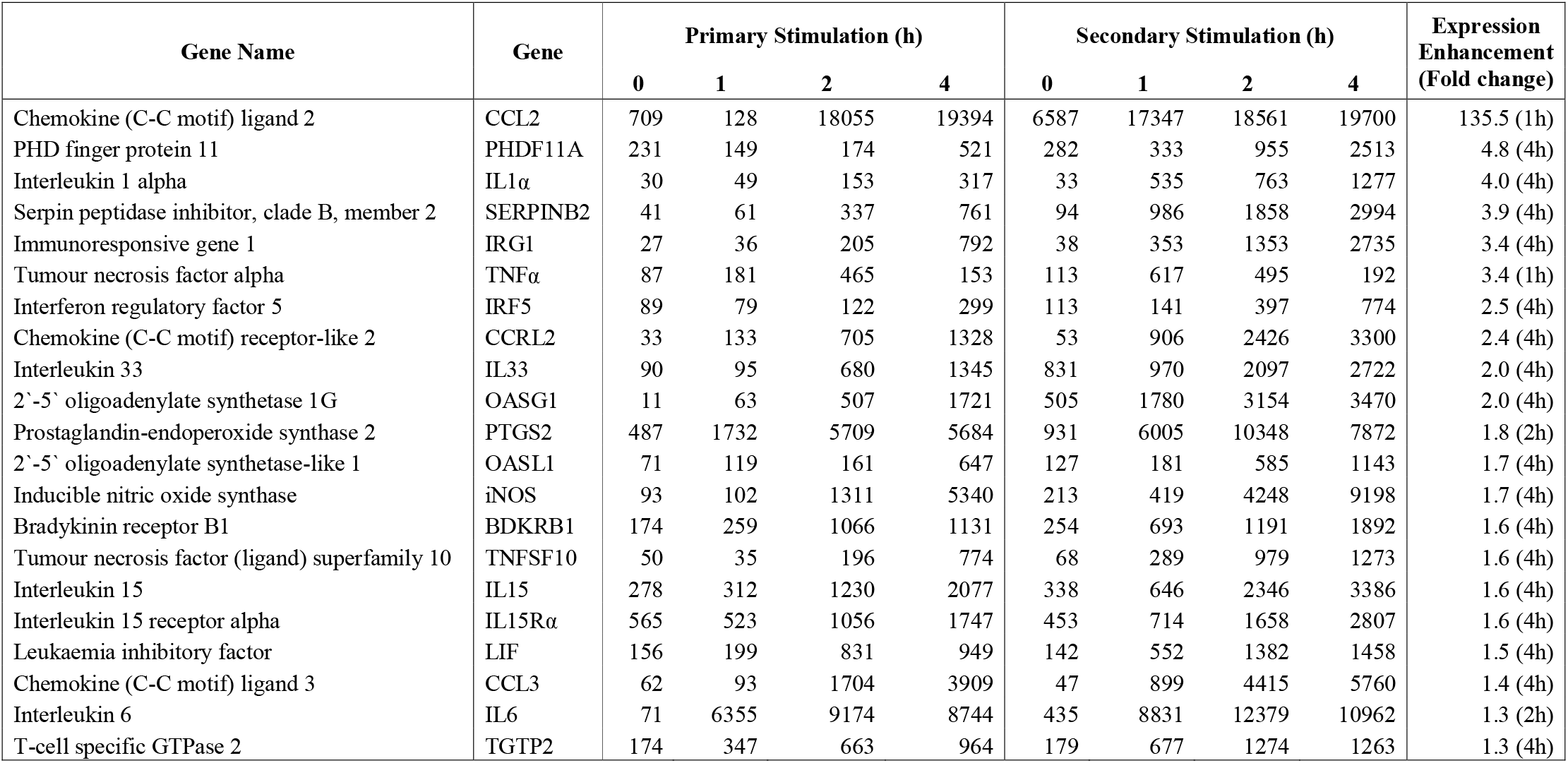
Genes that show transcriptional memory.

| Gene Name | Gene | Primary Stimulation (h) |  |  |  | Secondary Stimulation (h) |  |  |  | Expression Enhancement (Fold change) |
| --- | --- | --- | --- | --- | --- | --- | --- | --- | --- | --- |
|  |  | 0 | 1 | 2 | 4 | 0 | 1 | 2 | 4 |  |
| Chemokine (C-C motif) ligand 2 | CCL2 | 709 | 128 | 18055 | 19394 | 6587 | 17347 | 18561 | 19700 | 135.5 (1h) |
| PHD finger protein 11 | PHDF11A | 231 | 149 | 174 | 521 | 282 | 333 | 955 | 2513 | 4.8 (4h) |
| Interleukin 1 alpha | IL1α | 30 | 49 | 153 | 317 | 33 | 535 | 763 | 1277 | 4.0 (4h) |
| Serpin peptidase inhibitor, clade B, member 2 | SERPINB2 | 41 | 61 | 337 | 761 | 94 | 986 | 1858 | 2994 | 3.9 (4h) |
| Immunoresponsive gene 1 | IRG1 | 27 | 36 | 205 | 792 | 38 | 353 | 1353 | 2735 | 3.4 (4h) |
| Tumour necrosis factor alpha | TNFα | 87 | 181 | 465 | 153 | 113 | 617 | 495 | 192 | 3.4 (1h) |
| Interferon regulatory factor 5 | IRF5 | 89 | 79 | 122 | 299 | 113 | 141 | 397 | 774 | 2.5 (4h) |
| Chemokine (C-C motif) receptor-like 2 | CCRL2 | 33 | 133 | 705 | 1328 | 53 | 906 | 2426 | 3300 | 2.4 (4h) |
| Interleukin 33 | IL33 | 90 | 95 | 680 | 1345 | 831 | 970 | 2097 | 2722 | 2.0 (4h) |
| 2'-5' oligoadenylate synthetase 1G | OASG1 | 11 | 63 | 507 | 1721 | 505 | 1780 | 3154 | 3470 | 2.0 (4h) |
| Prostaglandin-endoperoxide synthase 2 | PTGS2 | 487 | 1732 | 5709 | 5684 | 931 | 6005 | 10348 | 7872 | 1.8 (2h) |
| 2'-5' oligoadenylate synthetase-like 1 | OASL1 | 71 | 119 | 161 | 647 | 127 | 181 | 585 | 1143 | 1.7 (4h) |
| Inducible nitric oxide synthase | iNOS | 93 | 102 | 1311 | 5340 | 213 | 419 | 4248 | 9198 | 1.7 (4h) |
| Bradykinin receptor B1 | BDKRB1 | 174 | 259 | 1066 | 1131 | 254 | 693 | 1191 | 1892 | 1.6 (4h) |
| Tumour necrosis factor (ligand) superfamily 10 | TNFSF10 | 50 | 35 | 196 | 774 | 68 | 289 | 979 | 1273 | 1.6 (4h) |
| Interleukin 15 | IL15 | 278 | 312 | 1230 | 2077 | 338 | 646 | 2346 | 3386 | 1.6 (4h) |
| Interleukin 15 receptor alpha | IL15Rα | 565 | 523 | 1056 | 1747 | 453 | 714 | 1658 | 2807 | 1.6 (4h) |
| Leukaemia inhibitory factor | LIF | 156 | 199 | 831 | 949 | 142 | 552 | 1382 | 1458 | 1.5 (4h) |
| Chemokine (C-C motif) ligand 3 | CCL3 | 62 | 93 | 1704 | 3909 | 47 | 899 | 4415 | 5760 | 1.4 (4h) |
| Interleukin 6 | IL6 | 71 | 6355 | 9174 | 8744 | 435 | 8831 | 12379 | 10962 | 1.3 (2h) |
| T-cell specific GTPase 2 | TGTP2 | 174 | 347 | 663 | 964 | 179 | 677 | 1274 | 1263 | 1.3 (4h) |

Whilst it was encouraging that iNOS and IL-6 were amongst the genes identified as displaying transcriptional memory in the arrays, only a single microarray was performed for each sample and so it was necessary to confirm these observations by more quantitative methods using a greater number of preparations. To this end, the experiment was repeated multiple times (as shown in the legends to individual tables and figures) and qPCR used to measure transcript levels. The behavior of all the targets measured (over half of those shown in Table 1) was verified using this approach. Figure 3 shows four representative genes (CCL2, IL-1a, TNF-a and PDHF11A). In general, the secondary response shows the same kinetics as the primary response, following the same pattern of increase and decay, but the rise in expression either occurs sooner or is more exaggerated in amplitude.

**Figure 3.**
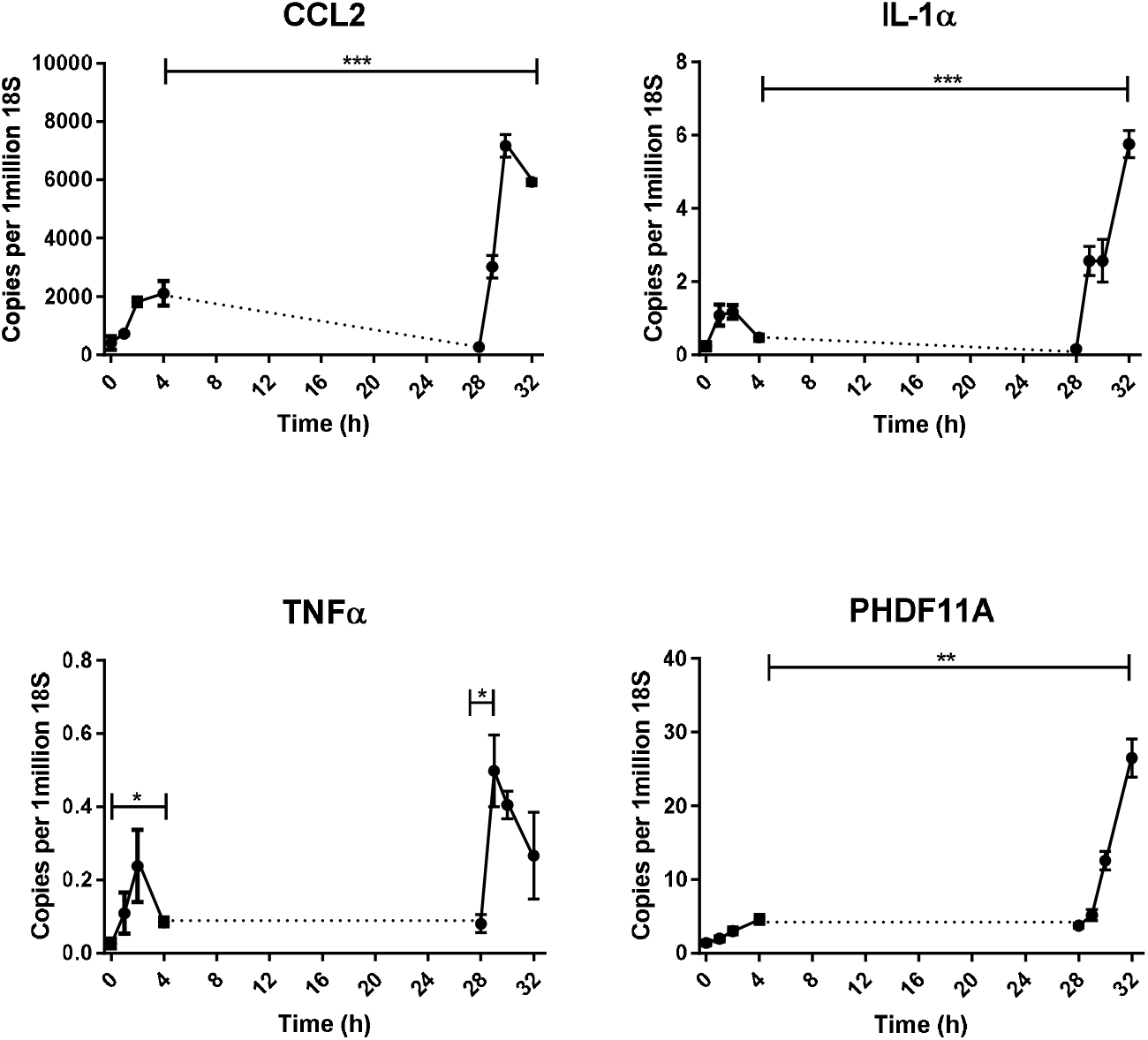
Expression of inflammatory genes in 3T3-L1 adipocytes during primary and secondary MCM exposures that show transcriptional memory. 3T3-L1 adipocytes were exposed to MCM for 4h (primary stimulation) and then exposed again for another 4h (secondary stimulation) after receiving a 24h washout in normal complete growth medium. cDNA was synthesised from RNA extracts and gene expression levels were analysed using qPCR. C_T_ results were normalised to 18S rRNA and expressed as transcript copies per 10^6^ 18S copies. Data are shown as mean ±SEM (*n*=3). Statistical analysis was performed using students t-test, p< 0.05*, <0.01**, <0.001***, <0.0001****

Microarray analysis revealed a further class of targets whose expression does not decrease after removal of the MCM stimulus (Table 2). For these genes, the transcript levels remain elevated (and often even keep increasing) even after 24 hours of washout. In most, but not all cases, expression levels increased even more on secondary exposure. As before, we confirmed these patterns using qPCR in new samples; Figure 4 shows data for one example: SAA3. As with the genes that showed transcriptional memory in Table 1, the genes that show persistence of gene expression elevation even after removal of the MCM stimulus all have at least one ontology which classifies them as inflammatory.

**Table 2.**
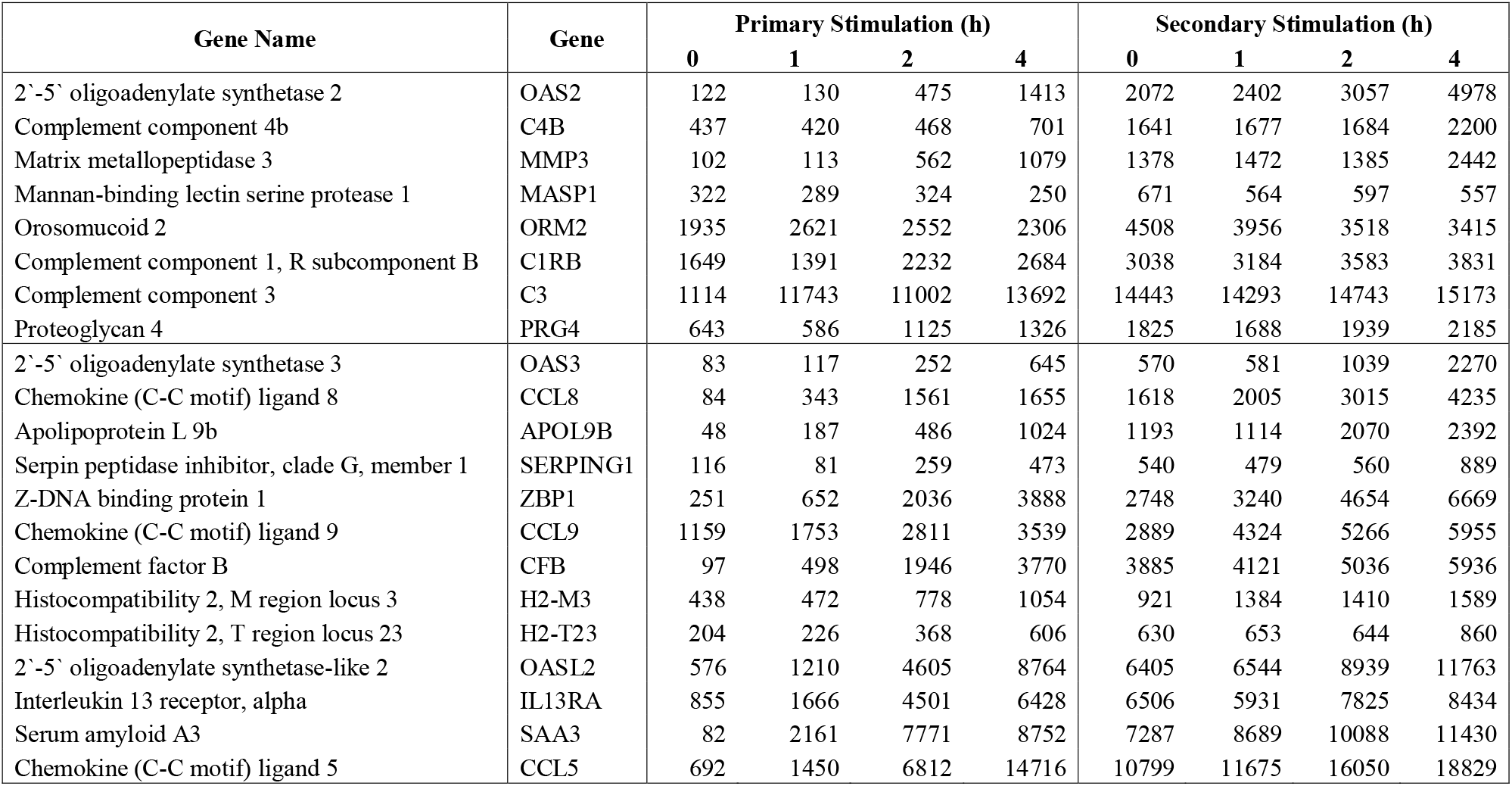
Genes with persistent expression after primary MCM stimulation.

**Figure 4.**
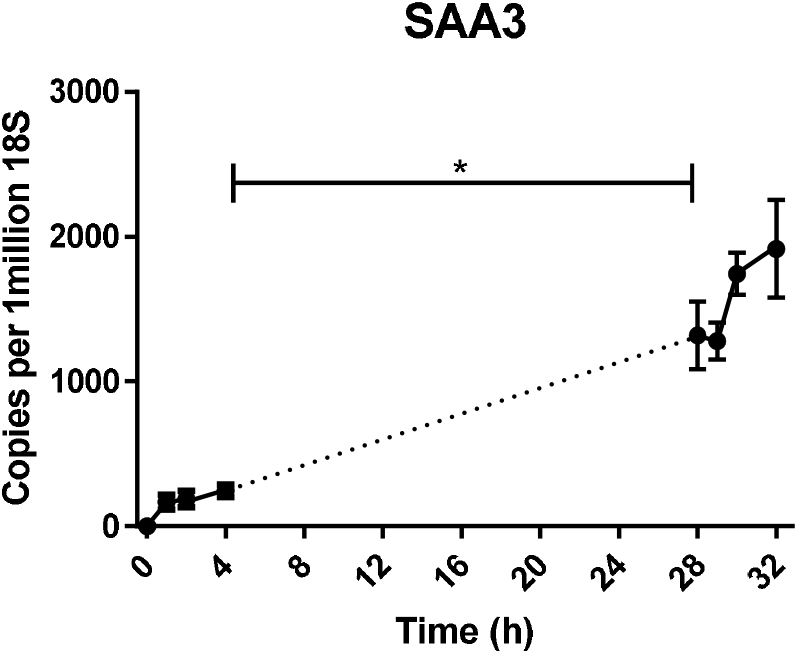
Expression of SAA3 in 3T3-L1 adipocytes during primary and secondary MCM exposures. 3T3-L1 adipocytes were exposed to MCM for 4h (primary stimulation) and then exposed again for another 4h (secondary stimulation) after receiving a 24h washout in normal complete growth medium. cDNA was synthesised from RNA extracts and gene expression levels were analysed using qPCR. C_T_ results were normalised to 18S rRNA and expressed as transcript copies per 10^6^ 18S copies. Data are shown as mean ±SEM (*n*=3). Statistical analysis was performed using students t-test, p< 0.05*, <0.01**

### Genes exhibiting Transcriptional Memory share only loose commonality in Promoter Properties

The promoters associated with the sequences in Table 1 and Table 2 were interrogated using the module in the Genomatix Software Suite (https://www.genomatix.de) that identifies common transcription factor binding sites. Analysis included all the members of Tables 1 (20 sequences) and Table 2 (21 sequences) except Masp1 for which information was not available.

There were 87 transcription factor binding sites (TFBs) that were common between at least 17 of the 20 genes in Table 1. Of these, a subset of 16 TFBs were characterised as being selectively concentrated in these genes in comparison to the mouse genome as a whole (P<0.0001). Those with the most extensive commonality included AP4R (transcription factor AP4), HUB1 (HTLV-I U5 repressive element-binding protein 1), GFI1 (Growth factor independence transcriptional repressor), IKRS (Ikaros zinc finger family), BHLH (bHLH transcription factors) and ZF12 (C2H2 zinc finger transcription factor family). Similarly, the promoters of the 20 genes from Table 2 showed a total of 64 types of TFB, of which 10 were identified as being distinct for this set. The three with the most penetration across the 20 genes were NRSF (Neuron-restrictive silencer factor), SF1F (Vertebrate steroidogenic factor) and STAF (Selenocysteine tRNA activating factor). There was some commonality in TFBs between the two tables, but not in any of the afore-mentioned genes.

### Genes repressed by MCM exposure do not show Transcriptional Memory

Exposure of adipocytes to MCM reduced the expression of several genes, and these are shown in Table 3, with selected qPCR confirmation from separate biological preparations shown in Figure 5. The horizontal divider in Table 3 delineates transcripts that recover to pre-stimulation levels after the washout from those that remain repressed after removal of the first stimulus. The targets in Table 3 are generally associated with ontological groupings related to adipocyte biology. Interestingly, even if there was full recovery of the transcript level during the washout period, the secondary response never showed an overtly faster or high amplitude response than the initial exposure.

**Table 3.** Genes with decreased expression upon MCM stimulation.

| Gene Name | Gene | Primary Stimulation (h) |  |  |  | Secondary Stimulation (h) |  |  |  |
| --- | --- | --- | --- | --- | --- | --- | --- | --- | --- |
|  |  | 0 | 1 | 2 | 4 | 0 | 1 | 2 | 4 |
| RAS, dexamethasone-induced 1 | RASD1 | 2214 | 330 | 243 | 460 | 1603 | 285 | 180 | 327 |
| Insulin receptor substrate 1 | IRS1 | 1427 | 1066 | 548 | 433 | 1164 | 1055 | 716 | 557 |
| CCAAT/enhancer-binding protein alpha | CEBPα | 1739 | 1628 | 932 | 574 | 1501 | 1149 | 962 | 616 |
| Lipin 1 | LPIN1 | 2906 | 3162 | 2014 | 1006 | 1726 | 1667 | 1416 | 829 |
| Low-density lipoprotein receptor-related protein 6 | LRP6 | 5850 | 5034 | 3848 | 2342 | 5204 | 4989 | 4102 | 2235 |
| Retinoic acid receptor, alpha | RARA | 808 | 729 | 513 | 375 | 1160 | 934 | 673 | 472 |
| Insulin receptor substrate 2 | IRS2 | 1315 | 2695 | 1904 | 619 | 1190 | 1083 | 667 | 709 |
| Peroxisomal biogenesis factor 11 alpha | PEX11A | 935 | 758 | 673 | 458 | 1179 | 987 | 721 | 527 |
| Peroxisome proliferator-activated receptor gamma | PPARγ | 7849 | 7766 | 5926 | 4476 | 8103 | 7823 | 7279 | 5442 |
| Laminin, alpha 4 | LAMA4 | 5465 | 4521 | 4416 | 3619 | 5026 | 5306 | 5061 | 4406 |
| Hormone-sensitive lipase | HSL | 1738 | 1752 | 1552 | 1201 | 1830 | 1795 | 1490 | 966 |
| Perilipin 4 | PLIN4 | 4427 | 4141 | 3778 | 3302 | 4564 | 4536 | 3495 | 2642 |
| Phosphoenolpyruvate carboxylase kinase 1 | PCK1 | 1923 | 1160 | 665 | 328 | 326 | 300 | 168 | 177 |
| Insulin-induced gene 1 | INSIG1 | 8718 | 9154 | 6401 | 4711 | 5500 | 5698 | 5388 | 3281 |
| Adipogenin | ADIG | 821 | 552 | 591 | 449 | 483 | 347 | 295 | 227 |
| Leptin | LEP | 155 | 132 | 113 | 91 | 85 | 69 | 77 | 53 |
| Glucose transporter type 4 | GLUT4 | 3889 | 3820 | 3472 | 2783 | 2476 | 2375 | 2138 | 1786 |
| Fatty acid synthase | FAS | 12293 | 12552 | 10305 | 9779 | 8274 | 8569 | 7367 | 5277 |
| Fatty acid binding protein 5 | FABP5 | 1116 | 1037 | 1055 | 963 | 897 | 824 | 770 | 673 |
| Lipoprotein lipase | LPL | 18124 | 16207 | 15929 | 16211 | 15839 | 15772 | 14698 | 13811 |
| Diglyceride O-acyltransferase 1 | DGAT1 | 6898 | 10381 | 10032 | 8948 | 8700 | 8118 | 7718 | 6771 |

**Figure 5.**
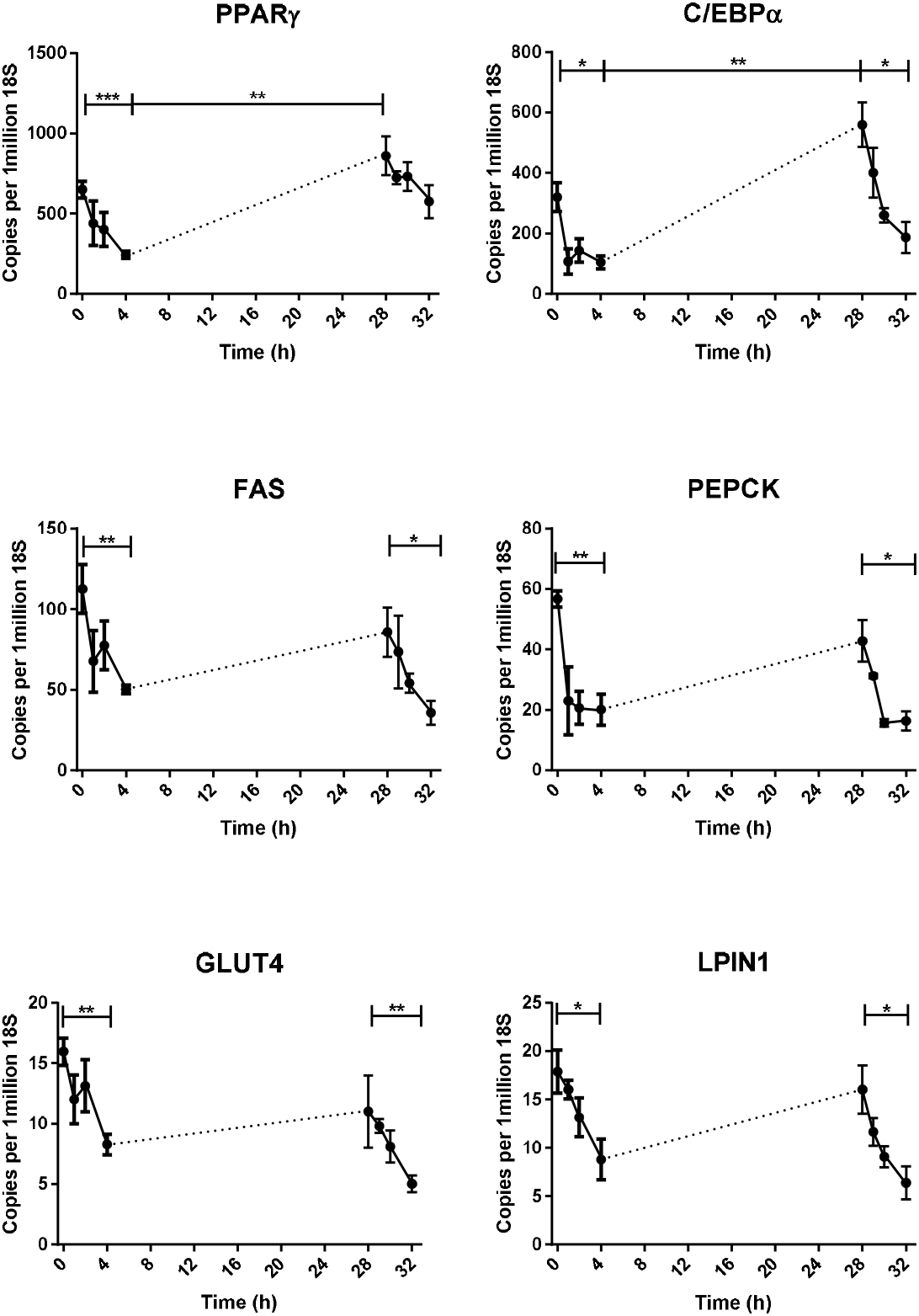
Expression of genes in 3T3-L1 adipocytes during primary and secondary MCM exposures. 3T3-L1 adipocytes were exposed to MCM for 4h (primary stimulation) and then exposed again for another 4h (secondary stimulation) after receiving a 24h washout in normal complete growth medium. cDNA was synthesised from RNA extracts and gene expression levels were analysed using qPCR. C_T_ results were normalised to 18S rRNA and expressed as transcript copies per 10^6^ 18S copies. Data are shown as mean ±SEM (*n*=3). Statistical analysis was performed using students t-test, p< 0.05*, <0.01**, <0.001***, <0.0001****

### Adipocytes retain but do not magnify the memory effect following multiple MCM challenges

Figure 6 shows the results of several rounds of MCM stimulation and washout on genes that show either an enhanced secondary response (iNOS) or persistent high expression after the first stimulus (SAA3). In the case of iNOS, the memory effect was retained after a second and third washout and so gave rise to accentuated responses (compared to the first exposure) on the third and fourth challenges. However, the extra exposures did not result in a greater level in expression relative to the second challenge. The first period of exposure and washout is therefore sufficient to elicit the full memory effect. In the case of SAA3, the first washout period was, as before, characterised by a continued rise in levels of this transcript, and the expression level remained highly elevated regardless of the removal or re-instigation of the stimulus.

**Figure 6.**
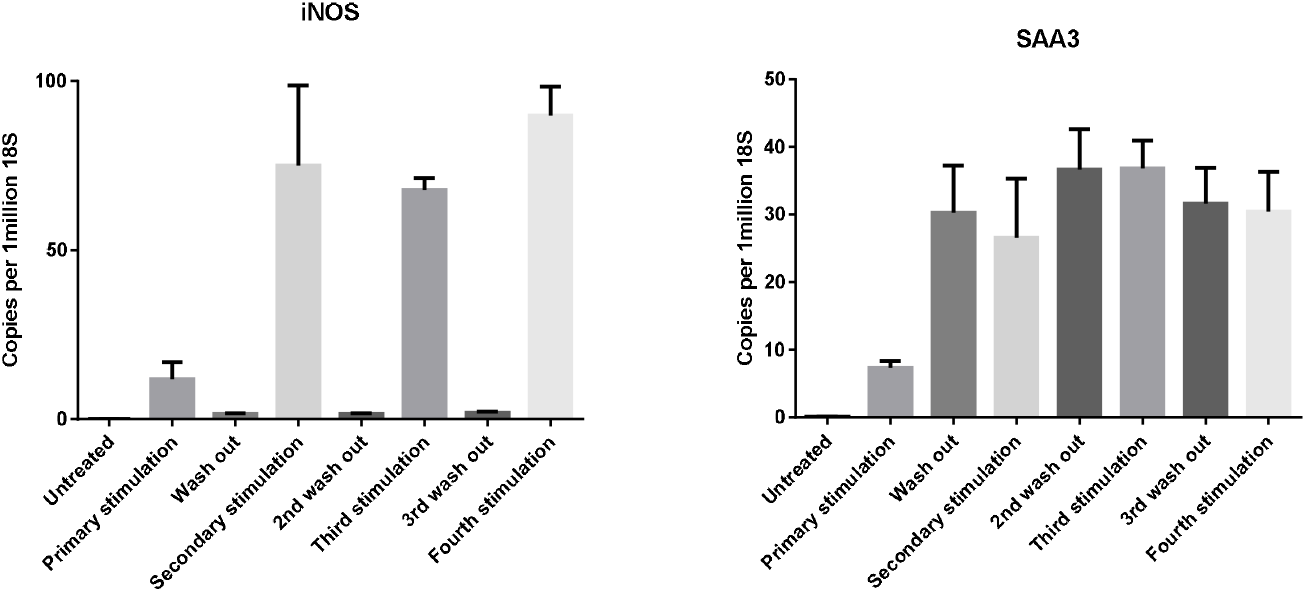
Expression of genes in 3T3-L1 adipocytes during primary, secondary, third and fourth MCM exposures. 3T3-L1 adipocytes were stimulated with MCM for 2h (primary) after receiving either one (secondary), two (third) or three (fourth) previous exposures with MCM for 4h followed by a 24h washout in normal growth medium. cDNA was synthesised from RNA extracts and gene expression levels were analysed using qPCR. C_T_ results were normalised to 18S rRNA and expressed as transcript copies per 10^6^ 18S copies. Data are shown as mean ±SEM (*n*=3). Statistical analysis was performed using students t-test, p< 0.05*, <0.01**, <0.001***

### Transcriptional memory is retained for at least one week

To determine the longevity of the memory effect after the initial stimulus, we extended the time of the washout period between MCM challenges to up to 12 days. Because the adipocytes were already 12-days post-differentiation at this stage, and because the transcriptome profile of 3T3-L1 cells varies over such a long period (32), age-matched naive cells were included as controls at each time point. Figure 7 shows the results of this experiment on iNOS expression. Because the level of this transcript in naive cells and in cells after washout is always close to undetectable, these values are omitted for clarity. As before, after 1 day of washout, there is a pronounced memory effect, with the secondary stimulus giving rise to a 3-fold greater response than the primary exposure. Despite the fact that the absolute magnitude of the MCM-stimulation decreased with cell age, a clear difference between the primary and secondary responses was still seen after three and six days post-washout of the initial stimulus. However, after a washout period of 12 days, the response of previously exposed cells was not different to that of naive (never-exposed) cells. It should be noted however, that the responsiveness of cells to MCM stimulus at this stage (some 24 days post-differentiation) was only about 50% of that of younger cells. Therefore, at least in the case of iNOS, the effects of the primary MCM exposure persist for at least six days post-washout.

**Figure 7.**
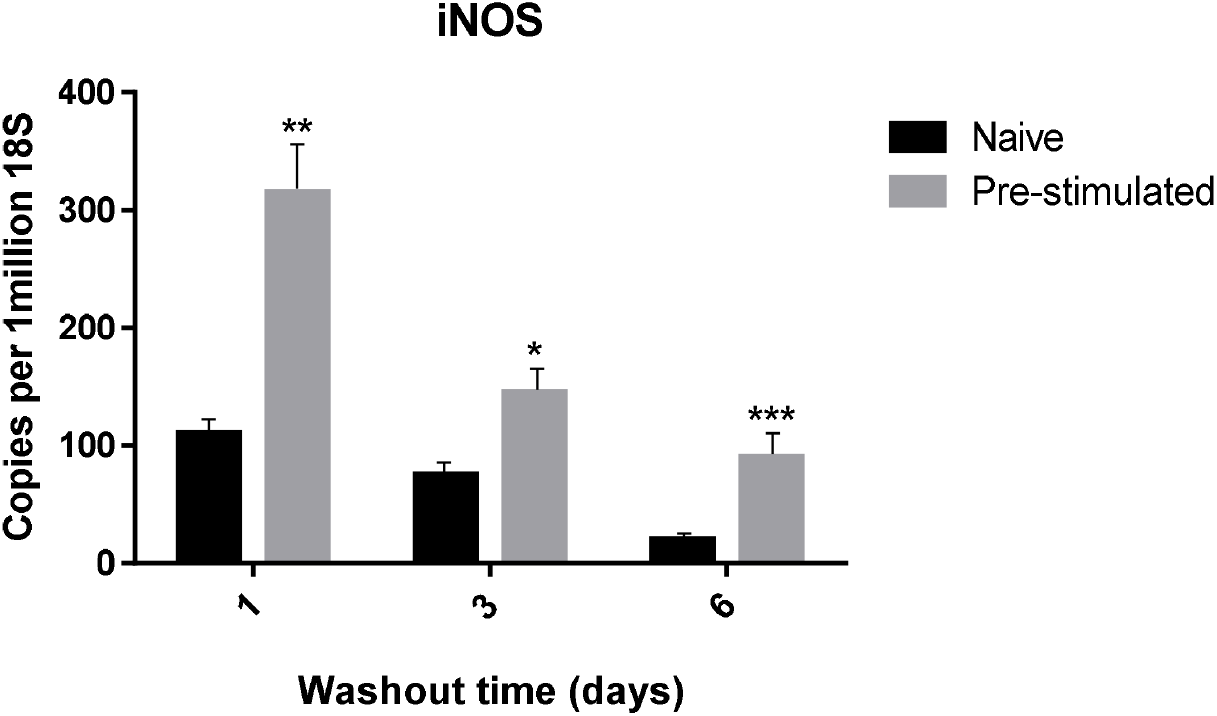
Expression of iNOS in 3T3-L1 adipocytes during primary and secondary MCM exposures with extended washout times. 3T3-L1 adipocytes were either grown in normal growth medium (naïve) or exposed to MCM for 4h (pre-stimulated) and then both cell groups were subsequently exposed to MCM for 4h after receiving a 1, 3, 6 or 12 day washout in normal complete growth medium. cDNA was synthesised from RNA extracts and gene expression levels were analysed using qPCR. C_T_ results were normalised to 18S rRNA and expressed as transcript copies per 10^6^ 18S copies. Data are shown as mean ±SEM (n=3). Statistical analysis was performed using students t-test, p< 0.05*, <0.01**, <0.001*** naïve compared to pre-stimulated.

### Transcriptional Memory does not involve a change in the sensitivity to MCM

In order to determine if the primary MCM exposure alters the sensitivity or responsiveness to subsequent MCM exposure, the dose-response relationship for naive and pre-stimulated cells was established (Figure 8). In both groups of cells, iNOS expression was not raised by MCM if it was diluted by more than 100-fold. However, a 1:50 dilution of MCM was sufficient to cause a change in the level of iNOS transcript in both groups. At higher concentrations of MCM, the iNOS expression was always greater in cells that had been pre-exposed to MCM. Whereas iNOS expression appeared to plateau when naive cells were incubated with undiluted MCM, in pre-exposed cells the level of iNOS expression appeared to be proportional to MCM concentration. Since more concentrated MCM was not available, it was not possible to determine the MCM concentration at which the response of pre-exposed cells would plateau. Regardless, these data show that the effect of prior MCM exposure is to increase the responsiveness of the iNOS expression system to subsequent MCM exposure, rather than increasing the sensitivity of the process.

**Figure 8.**
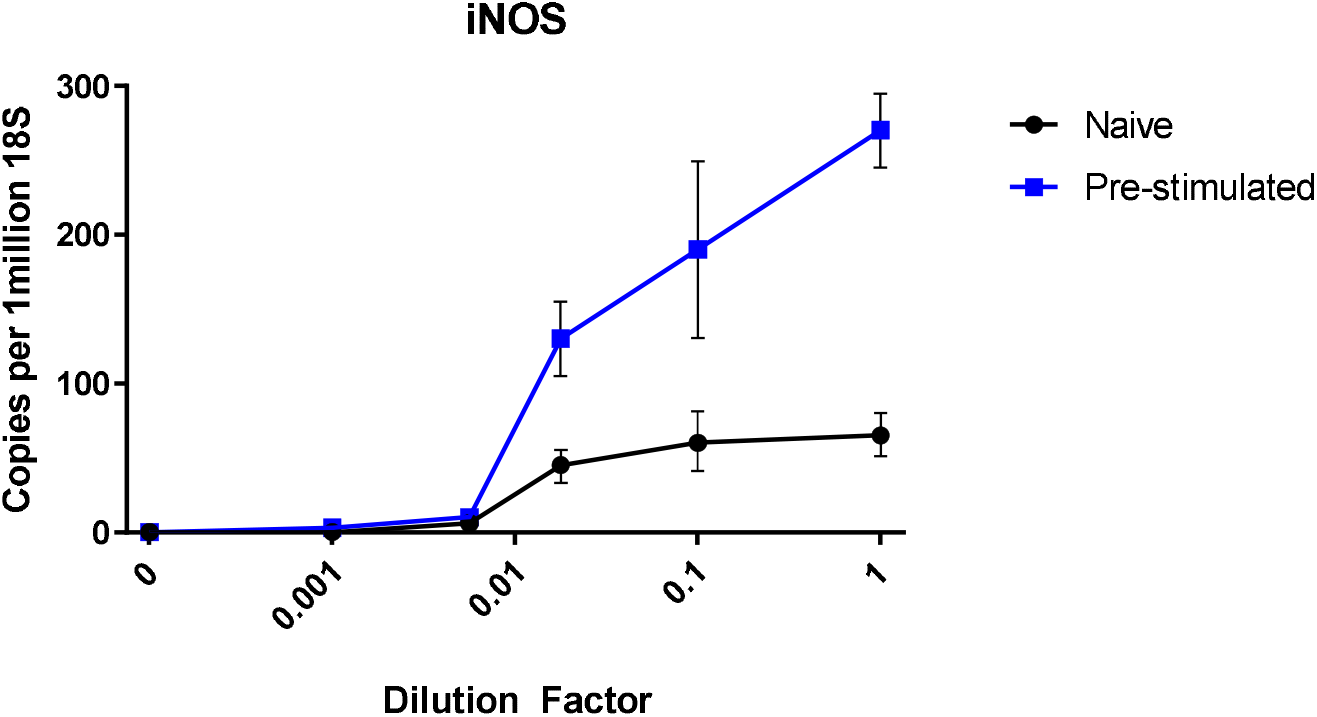
Expression of iNOS in 3T3-L1 cells during primary and secondary stimulation with varying concentrations of MCM. 3T3-L1 adipocytes were either grown in normal growth medium (naïve) or exposed to 1:1 MCM for 4h (pre-stimulated) and then both cell groups were subsequently exposed to varying concentrations of MCM for 4h after receiving a 24h washout in normal complete growth medium. cDNA was synthesised from RNA extracts and gene expression levels were analysed using qPCR. C_T_ results were normalised to 18S rRNA and expressed as transcript copies per 10^6^ 18S copies. Data are shown as mean ±SEM (n=3). Statistical analysis was performed using students t-test, p< 0.05*, <0.01**

## DISCUSSION

Recent research (33) (34) (35) (36) (37) (38) suggests that cells of the innate immune system exhibit a form of transcriptional memory in that a priming event alters the cells so that later re-exposure to a similar stimulus causes an exaggerated response. Our results indicate that adipocytes possess a similar transcriptional memory to macrophage secretions and that it is manifested in two ways. Firstly, elevated expression of some genes persists long after removal of the stimulus. Secondly, and more intriguingly, the response of a subset of genes is reproducibly enhanced following a second challenge, despite the fact that their expression levels return to baseline between exposures. It is important to appreciate that although we have reported detailed data for a limited number of targets (e.g., iNOS, IL-6, SAA3, etc.) we have confirmed the existence of the memory response for more than 10 of the genes identified by microarray analyses.

In trying to dissect out the molecular mechanisms that might underlie this phenomenon, it is important to reflect on the nature of the stimulus used, and the normal processes that operate to control the waxing and waning of the resulting transcriptional response. It is likely that most of the stimulants in the MCM are cytokines and residual lipopolysaccharide. After an initial surge in expression, caused by stimulation of genes such as Toll-like receptors and interleukin receptors, braking mechanisms are rapidly initiated (for example the concomitant activation of the counter-regulatory SOCS pathway (39) as recently reviewed by (17). Rapid clearance of the initially produced transcripts is aided by the fact that most of the responsive mRNAs have a very short half-life. Therefore, the ‘transcriptional memory’ may be a consequence of increased initiation factor or RNA polymerase activity, an impaired dampening response or even a change in the turnover of transcripts.

We do know that a single exposure to MCM is sufficient to prime the memory, and that multiple cycles of washout and re-exposure do not give rise to lesser or greater memory effects. Another important clue is that, in the case of genes like iNOS, after each cycle of washout, the expression of every memory-enabled transcript returns to basal levels. Therefore, whatever imprint on the transcriptional system is responsible for the memory requires a second stimulus for it to be revealed. In contrast, in the case of genes like SAA3, a single exposure appears to render the transcriptional apparatus continually active, even in the absence of the stimulus. Significantly, it does not increase further upon subsequent stimulation, which implies that the initial exposure is sufficient to set in train transcriptional processes that are hard to switch off.

We did not delineate precisely which components of MCM are required to give rise to the transcriptional memory phenomenon. MCM will comprise of tens, if not hundreds, of inflammogens and, furthermore, although always prepared and used in the same way, each preparation of MCM will likely have had a slightly different composition. We were, however, able to establish that the LPS residual in MCM preparations was not responsible for the memory effect. Thus, although LPS is effective at raising the expression of genes like iNOS and IL-6, it does so at a much lower magnitude than MCM and subsequent cycles of washout and re-exposure do not result in a different response (results not shown).

It was established that the primary exposure does not increase the sensitivity of adipocytes to MCM but, rather, the total capacity of the inflammatory gene response seems to be accentuated by a single four-hour exposure to MCM. Changes in responsiveness are usually the result of a physical increase in the number of components in a signaling system – either at the receptor level (40) (41) (42), the intermediate signaling molecules (43) (12) (44) (45) or, more usually, in the total amount of the final target (46) (47). In the context of the current system, the final target could be envisaged to be active RNA polymerase bound to the promoter region of the genes. It is becoming increasingly clear that initiation of transcription is only partly responsible for the regulation of the rate of transcript production (48). After the assembly of RNA polymerase and relevant transcription factors on open chromatin in the promoter area, the efficiency of elongation can be differentially regulated, and this is especially true of genes that are highly regulated (49). Indeed, it is possible to convert a relatively unstable transcription complex into a unit with enhanced processivity (50). In other cases in which there are enhanced secondary responses, termination of the primary stimulus leaves the RNA polymerase still associated with the activated genes. Thus the RNA polymerase is poised, ready for the arrival of the next stimulus (51). Although we have not directly measured the location of RNA polymerase after the first stimulus, our data is entirely consistent with this mechanism operating for the targets that show TM. Clearly, measuring the presence of RNA pol II around the promoter area will be crucial in determining if the first stimulus leaves RNA pol II poised in this manner.

At a more general level, if the chromatin around the promoter region of a gene is open then it is easier for gene expression to be initiated than if the nucleosomes in same area of chromatin are highly organized and densely packed. The observations that the first exposure seems to increase the total responsiveness of the system and that multiple exposures do not incrementally increase the TM effect, are both consistent with the primary exposure simply making the chromatin maximally accessible. Techniques now exist to determine the extent to which particular sections of chromatin are accessible (e.g., FAIRE (52) (53) (54)and mono nuclease digestion (55) (52) (56)) and these approaches will be important in trying to elucidate the molecular mechanisms involved. Although we were not successful in identifying specific promoter modules that might be responsible for the two types of TM, there was enough consistency in the groups to inform the choice of probes in future experiments aimed at determining if particular transcription factors are involved. However, although the nearly 60 transcription factor modules present on the genes in question also appear with high frequency on all inflammatory genes, the fact that not all inflammatory genes show the memory response indicates that other factors must be involved. Accordingly, and consistent with the memory in the innate immune response, the priming event may lay down epigenetic markers in particular areas of the genome and it is also possible that long non-coding RNAs and micro-RNAs are involved. These are obviously areas for further investigation.

Another mechanistic clue may be in the fact that the memory effect is preserved for at least six days after the initial exposure, especially as this is despite the inclination of differentiated 3T3-L1 adipocytes to become less sensitive to MCM with increasing time in culture. Therefore whatever mechanisms operate to reduce MCM responsiveness during extended culture are not abrogated by an previous MCM exposure but, equally, the mechanisms responsible for the memory effect are clearly still operational and can impose themselves in long-term cultured cells.

The ability of MCM to reduce the expression of adipocyte-characteristic genes was of interest and, in particular, it was noted that targets coding for enzymes with a critical role in lipogenesis (eg, ACC and PEPCK) were not only reduced in expression by MCM but largely failed to recover during the 24 hour washout period, even undergoing further reductions in expression level on subsequent MCM exposure. Commensurate with this, we have observed that prolonged incubation of mature 3T3-L1 cells with MCM does elicit substantial de-differentiation and blunts the differentiation process (results not shown), and this is supported by other studies that implicate MCM in reducing markers of the adipocyte phenotype (57) (58) (59) (60) (61) (7) (8) (62).

The existence of adipocyte transcriptional memory introduces an exciting complexity into our understanding of the relationship between adipocytes and macrophages. Although this behavior should be confirmed *in vivo* before extrapolating the potentially far-reaching physiological consequences, it does emphasize the importance of better understanding the mechanisms which exist in adipose tissue to prevent an escalating positive feedback loop between the adipocyte and macrophage inflammatory secretions. If proven to be physiologically relevant, the phenomenon has implications for long-term health in diet cycles.

## Acknowledgements

We gratefully acknowledge the encouragement of Professor Joel Mackay in this research and for his criticism of the initial draft of the manuscript.

